# Spider webs as eDNA tool for biodiversity assessment of life’s domains

**DOI:** 10.1101/2020.07.18.209999

**Authors:** Matjaž Gregorič, Denis Kutnjak, Katarina Bačnik, Cene Gostinčar, Anja Pecman, Maja Ravnikar, Matjaž Kuntner

## Abstract

The concept of environmental DNA (eDNA) utilizes nucleic acids of organisms directly from the environment. Recent breakthrough studies have successfully detected a wide spectrum of prokaryotic and eukaryotic eDNA from a variety of environments, ranging from ancient to modern, and from terrestrial to aquatic. These numerous sources promise to establish eDNA as a tool for diverse scientific settings. Here, we propose and establish spider webs as a source of eDNA with far reaching implications. First, we conducted a field study to track specific arthropod targets from different spider webs. We then employed high-throughput amplicon sequencing of taxonomic barcodes to investigate the utility of spider web eDNA for biodiversity monitoring of animals, fungi and bacteria. Our results show that genetic remains on spider webs allow the detection of even the smallest target organisms. We also demonstrate that eDNA from spider webs is useful in research of community compositions in different domains of life, with potentially highly detailed temporal and spatial information.

## Introduction

The continuous loss of biodiversity is among the critical societal challenges^1–3^. Biodiversity data, crucial to countering these trends, still mostly rely on traditional taxonomic species identification, such as morphological and behavioral species-diagnostic traits. These traditional methods have well-known shortcomings, e.g. the majority of biodiversity is undescribed^4,5^, phenotypic diagnoses are often insufficient^6^, and crucial taxonomic expertise is declining^7,8^. As a result, traditional approaches to conducting efficient and standardized biodiversity surveys are often inadequate. In addition, traditional sampling techniques are invasive. They are likely to kill the target organisms and may further harm their ecosystems^3,9^.

Novel molecular approaches, such as sampling of environmental DNA (eDNA), i.e. obtaining intra- or extracellular genetic material directly from environmental samples, has the potential to overcome many limitations of traditional biodiversity monitoring^3,10–12^. eDNA is present in all media, e.g. soil, sediment, water and permafrost, and depending on the medium, the preservation of eDNA varies from a few weeks to hundreds of thousands of years^3,13,14^. eDNA can be detected either using targeted tests within a single-species approach, or using generic high-throughput sequencing in a multiple-species approach, e.g., by DNA metabarcoding^15^. Accordingly, eDNA is increasingly used to address fundamental questions in basic and applied research fields such as ecology, molecular biology, nature conservation, and paleontology^3,11,16–18^. Recent studies using eDNA have assessed plant^18,19^, fungal^20,21^, earthworm^22^, amphibian^23^ and fish^24^ communities, characterized the functional potential of microbial and viral communities^25^, monitored invasive species^52,53^, and detected difficult-to-find species such as cave olms^26^, brown bears^27^ and whale sharks^28^.

In this paper, we argue that spider webs represent a powerful tool for obtaining eDNA. Spiders are among the dominant predators of arthropod communities^29^. Not all spiders construct capture webs, but those that do, show enormous diversity and ecological abundance. Their webs range from millimeters to meters in size, are diverse in architecture, in the type of silk they consist of, and in the microhabitat they are suspended in^19,20^. Spider webs are ubiquitous in both natural and anthropogenic ecosystems. Their potential use as a source of eDNA has been proposed only recently. One pioneering study in controlled laboratory conditions demonstrated that widow spider webs contain genetic traces of the host and its single prey^30^. Another preliminary study confirmed these findings by amplifying spider DNA from silk of two host species^31^. Recently, a metabarcoding study demonstrated that spider webs are passive aerial filters, and thus potentially represent a useful new source of eDNA^32^. However, these early studies have limitations (see Supplementary material 1) and we believe that the utility of eDNA from spider webs far surpasses simple identification of spiders and their prey.

To move towards a methodology that utilizes spider webs as a source of eDNA, we need to advance our understanding of two critical issues. The first revolves around a single species detection in a web. This is achieved by testing the performance of species-specific detection of eDNA from spider webs over a wide dynamic range of concentrations of the target’s DNA, and including the establishment of appropriate DNA isolation and amplification controls. The second issue is whether webs, as aerial filters, can be used to identify multiple organisms. Given that temporal and spatial factors affect spider web biology, comparative metabarcoding should yield sufficient variation in species composition, as well as abundances, for biodiversity estimations. For example, many spider species build webs in specific microhabitats, which could provide fine spatial information. Also, orb web spiders completely renew their webs daily, while webs of other spider families are long-lasting^33^, which can provide useful temporal information. Furthermore, some but not all spider webs contain capture threads that are coated in viscous glue (e.g. orb webs)^34^, and could thus function differently in accumulating genetic material compared to webs that only contain “bare” silk threads (e.g. sheet webs).

Here, we employ the single- and multi-species approaches to estimate the utility of eDNA from spider webs. Within our single-species approach, we conduct field tests and apply rigorous laboratory protocols to test whether model prey can be detected from spider webs. These tests use quantitative PCR (qPCR) to detect prey DNA in the webs of the garden spider *Araneus diadematus* and the common hammock-weaver *Linyphia triangularis,* two species with different web types. Our multi-species approach then establishes metabarcoding protocols. In order to investigate how eDNA from spider webs may be used for biodiversity monitoring, we sampled webs of the above spider species in two distinct forest types and in two subsequent years. The detected animal, fungal, and bacterial diversities strongly support our prediction that eDNA accumulated in spider webs contains important biodiversity information.

## Methods

### Single-species eDNA detection

In a forest in Slovenia, we selected five webs of both the garden spider *Araneus diadematus* and the common hammock-weaver *Linyphia triangularis*. The garden spider builds a typical two-dimensional orb web, consisting of non-sticky, as well as sticky threads^34^. The viscous glue of these sticky capture threads is among the best bio-adhesives known^35^, likely making orb webs efficient in capturing parts of impacting arthropods and airborne particles. The common hammock-weaver builds a three-dimensional web, consisting of a hammock-like segment, interlaced with a looser mesh of silk above and below it. These webs do not contain sticky capture threads^34^. We introduced two small sized house crickets *Acheta domestica* (~ 5 mg body mass) into each web. We then collected the webs several hours later. The collected silk contained no visible prey remains. We collected additional five webs of each spider species as control samples. We collected each web onto the tip of a sterile plastic inoculation loop, breaking the loop off into a sterile microcentrifuge tube immediately after collection. For further processing, we stored all web samples at −80°C.

We isolated DNA from all samples using an adapted protocol of the PowerLyzer PowerSoil DNA Isolation Kit (Qiagen, USA; Supplementary material 2). We used quantitative (q)PCR that enables sensitive detection and quantification compared to regular PCR. In order to test the detection of house crickets in web samples, we designed and tested two TaqMan chemistry based assays targeting 102 bp and 127 bp fragments of the mitochondrial gene cytochrome c oxidase subunit I, respectively (COI; Supplementary material 2, Table S1). We provide amplification protocols in Supplementary material 2.

Throughout laboratory work, we employed rigorous protocol controls (Table 1): a negative isolation control, a positive amplification control, no template control during qPCR amplification, and an internal control of the DNA isolation process (using the “Eukaryotic 18S rRNA Endogenous Control”, Applied Biosystems, USA). To control for the performance of the two designed assays for detection of *A. domestica*, we employed further controls that we performed on two web types, using laboratory populations of the orb web building African hermit spider (*Nephilingis cruentata*) and the cobweb building Mediterranean black widow (*Latrodectus tredecimguttatus*). We tested the assays for DNA isolated from webs only (negative control) and from target (house cricket) tissue only (positive control). We employed assay specificity controls by testing the amplification of the host (spider) and non-target prey (mealworms) DNA. To control for potential inhibitors in the amplification matrix (spider silk), we tested the assays on samples of target tissue with added spider silk. To test the dynamic range of the assays, we tested both assays on a range of target DNA concentrations, by performing a dilution series of our test samples’ (webs of both species that were fed with house cricket) DNA in nuclease free water, from 10^−1^ to 10^−9^.

**Table 1:** Controls used for testing the performance of the qPCR assays used for the detection of *A. domestica*.

| Sample type | Control type | Adom1 (Cq) |  | Adom2 (Cq) |  |
| --- | --- | --- | --- | --- | --- |
|  |  | Undil. | 10x dil. | Undil. | 10x dil. |
| Cobweb, no target | Negative, primer specificity | neg. | neg. | neg. | neg. |
| Orb web, no target | Negative, primer specificity | neg. | neg. | neg. | neg. |
| Web host tissue (widow spider) | Primer specificity | neg. | neg. | neg. | neg. |
| Web host tissue (hermit spider) | Primer specificity | neg. | neg. | neg. | neg. |
| Laboratory prey (mealworm) | Primer specificity | neg. | neg. | neg. | neg. |
| Target (house cricket) | Primer specificity | 11.96 | 18.06 | 12.99 | 19.56 |
| Fresh cobweb + target tissue | Amplification inhibition | 12.31 | 18.87 | 13.30 | 20.39 |
| Fresh orb web + target tissue | Amplification inhibition | 15.09 | 22.01 | 16.33 | 23.49 |
| Cobweb from spider fed with target | Positive | 23.88 | 29.73 | 25.88 | 31.50 |
| Orb web from spider fed with target | Positive | 17.35 | 23.93 | 18.73 | 25.69 |

We considered a sample positive if it produced an exponential amplification curve that was distinguishable from the negative controls. In such cases, we determined the quantification cycles (Cq). For fluorescence acquisition and determination of Cq, we used the SDS 2.4 software (Applied Biosystems, USA). For this, we set the baseline between the 3rd and the 8th amplification cycle, and we set the fluorescence threshold manually at 0.05, i.e. at a level that was above the baseline and sufficiently low to be within the exponential increase region of the amplification curve.

### DNA metabarcoding

We sampled two forests in Slovenia from different climates, one submediterranean, the other continental. In both, the above described spider species *A. diadematus* and *L. triangularis* co-occur in close proximity. In each forest, we collected five webs of each species, as described in the “Single-species eDNA detection”. In the submediterranean forest, we repeated the sampling of *L. triangularis* webs in two subsequent years (25 samples in total, see Supplementary material 2, Table S6).

We isolated DNA from all samples using an adapted protocol of the PowerLyzer PowerSoil DNA Isolation Kit (Qiagen, USA; Supplementary material 2). As isolations for all web samples were successful, we randomly chose five web samples per web type per forest per year (25 samples total) for amplification. We amplified DNA from samples using primers commonly used in metabarcoding studies of arthropods, fungi and bacteria. For amplification of animal DNA, we used the primer set mlCOIintF and jgHCO2198 that target a 313 bp fragment of the mitochondrial gene cytochrome c oxidase subunit I (COI)^36^. For amplification of fungal DNA, we used the primer set ITS3_KYO2 and ITS4 that target a 300-400 bp fragment of the Internal transcribed spacer (ITS)^37^. For amplification of bacterial DNA, we used the primer pair S-D-Bact-0341-b-S-17 and S-D-Bact-0785-a-A-21 that targets a 464 bp fragment of 16S ribosomal RNA (16S)^38^. We provide other primer details and the amplification protocol in Supplementary material 2.

For all samples, we performed high-throughput sequencing of the amplicons using the Illumina MiSeq sequencing platform, through a commercial provider. Before sample submission, we measured the amount of amplified DNA for all samples, using microfluidic capillary electrophoresis on the Labchip GX (PerkinElmer), where we used the DNA High Sensitivity Assay. Based on these DNA amplicon amounts, we sequenced bacterial amplicons separately, as the amount of amplified bacterial DNA was several-fold larger compared to that of fungi and animals. For sequencing, we pooled amplified DNA of fungi and animals in a ratio of 3:1, as we expected more fungal OTUs compared to animal ones.

In all parts of sample preparation and sequencing, we included four negative control samples, obtained from individual DNA isolation procedures, and a microbial mock community sample (“ZymoBIOMICS Microbial Community Standard”, Zymo Research, Germany) that we used as a positive control in the metabarcoding experiment.

We analyzed sequence data using the QIIME2 2019.4 software package (Quantitative Insights Into Microbial Ecology)^39^. We trimmed primers, adapters, and regions with the quality score <20. Since the assembly of the paired end reads was not optimal, we used only single-end forward reads for analysis. We denoised the reads using DADA2, aligned representative sequences using mafft, and constructed the midpoint-rooted tree using FastTree. We calculated alpha and beta diversity indices. We used trained feature classifier to assign the sequences to taxonomic categories using three databases: Silva release 132 (bacteria)^40^, dynamically clustered UNITE ITS database version 8.0 (fungi)^41^, and BOLD downloaded on 31. 7. 2019 (animals)^42^. Since BOLD database is not provided in a format directly useful in QIIME2 and downloading of the complete database was not possible, we downloaded the fasta and tsv files using custom scripts for 238 smaller taxonomic groups, which contained sufficiently few data to result in successful download and later concatenation into a complete database covering all taxa present in BOLD. We concatenated these files and retained only sequences with ‘COI-3P’ or ‘COI-5P’ in the name. We replaced all characters not in the IUPAC nucleotide code with Ns. To produce the final set of sequences for use in QIIME2, we dereplicated the database with ‘vsearch’ using a threshold of 0.97^43^. We used alignment to the ITS and COI databases to separate ITS and COI sequences, which were pooled before sequencing – we discarded all non-aligning sequences (at the 90% cutoffs for the aligning parts of the sequence and the total sequence) using the option ‘qiime quality-control exclude-seqs’. Due to the large number of non-animal DNA in the COI sequences, we manually identified the representative sequences by blasting them against all barcode records on the BOLD database. We then manually removed all non-animal representative sequences from the QIIME2 representative sequences file, which was finally used to filter the feature table with the option ‘qiime feature-table filter-features’.

To render our samples comparable for diversity analyses, we randomly sampled reads from each sample and thus equalized their size. We retained 15900 reads for bacterial, 1360 reads for fungal, and 490 reads for animal samples, maximizing the number of samples rather than the number of reads (Supplementary material 2, Table S7). To investigate alpha diversity, we calculated the Chao1 and Shannon indices. We assessed differences in alpha diversity compositions using the Kruskal-Wallis H test. We determined the distance and dissimilarity matrices, to visualize the ordination and clustering of the bacterial, fungal and animal community compositions for beta diversity analyses, through weighted and unweighted Unifrac distance metrics^44^, and with the Jaccard similarity coefficient and Bray–Curtis dissimilarity. We evaluated the ordination patterns based on phylogenetic distance metrics using principal coordinate analysis (PCoA). We assessed differences in community compositions between sample types by non-parametric permutational analysis of variance (PERMANOVA), using the unweighted UniFrac distance metric and other settings at default within the QIIME2 pipeline.

## Results

To test the performance of the two assays designed for detection of *A. domestica*, we employed negative controls, primer specificity controls, and amplification controls. Both primer sets were positive for the house cricket (Table 1). In the dilution series, both primer sets showed consistent detection of a wide range of target DNA concentrations, with Adom1 primers consistently outperforming the Adom2 primers, i.e. having lower Cq values for the same samples (Fig. 1a, Table 1). Therefore, we used Adom1 in field tests. We successfully traced house crickets in all 10 webs, with Cq values ranging from 14 to 32 (Fig. 1b). We also successfully traced house crickets in all 10 samples when diluted 10-fold. All negative controls were negative, while the internal isolation control (18S assay) was positive in all samples.

**Figure 1:**
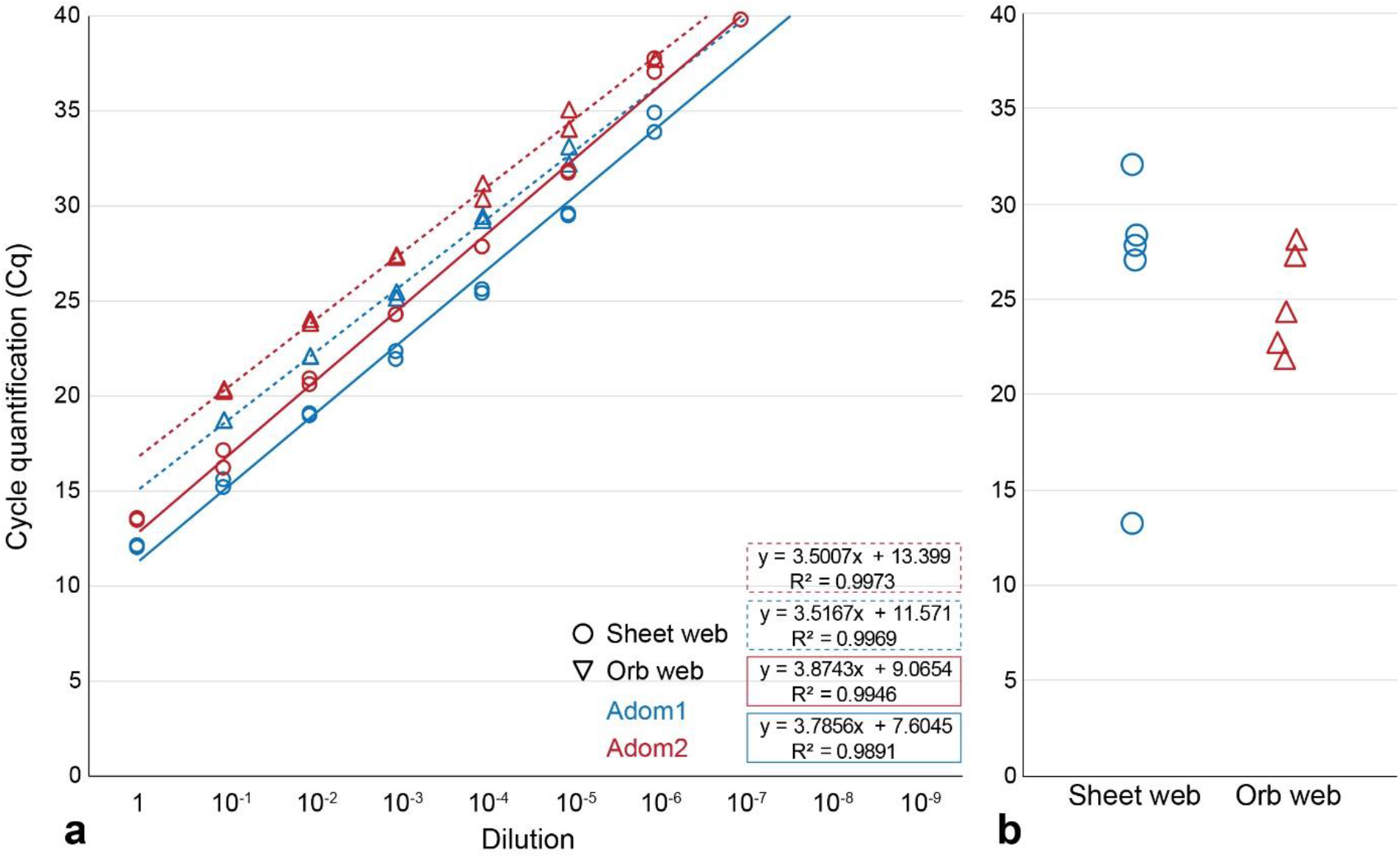
Targeted detection of house crickets from webs in nature. a) The dilution series of DNA isolated from two web types (designated by architecture) containing traces of prey sample (*Acheta domestica)* performed usingthe Adom1 and Adom2 qPCR assays (designated by color). a) Detection success of the target, showing the Cq values for Adom1 for samples of two web types containing traces of *A. domestica* DNA.

Our metabarcoding investigation retrieved a total of 2,164,380 bacterial reads, 721,594 fungal reads, and 174,006 animal reads from the 25 web samples. Summing up to 11,285 OTUs, bacteria represented the largest diversity, followed by fungi with 4005 OTUs, and animals with 314 OTUs (Supplementary material 3, Tables S1, S2, S4). Among bacteria, we identified 30 phyla (Supplementary material 3, Table S1). Among fungi, we identified 9 phyla, containing 31 classes, 108 orders and 307 families. Ascomycota and Basidiomycota were by far the most diverse, encompassing 297 of these 307 families (Supplementary material 3, Table S3). Among animals, we identified 3 phyla, containing 5 classes, 16 orders and 50 families. Insects were the most diverse, representing 33 of these 50 families (Supplementary material 3, Table S5). The microbial mock community sample that we used as control contained eight bacterial and two fungal species, of which we detected all but one bacterial species.

The statistical analysis of the chao1 index of alpha diversity showed that in the two forests, webs accumulate an equal diversity of bacterial (p = 0.950, H = 0.004), fungal (p = 0.412, H = 0.672) and animal (p = 0.143, H = 2.148) eDNA (Fig. 2). The sampling year yielded a similar eDNA diversity of the three organismal groups (bacteria: p = 0.314, H = 1.016; fungi: p = 0.456, H = 0.556; animals: p = 0.207, H = 1.595). Compared to orb webs, sheet webs accumulated a higher diversity of bacterial (p = 0.014, H = 6.036) and fungal (p = 0.022, H = 5.270), but not animal (p = 0.518, H = 0.418) eDNA. The statistical analysis of the Shannon index of alpha diversity showed similar results (Supplementary material 3). The described diversity pattern was reflected in the average OTU number per web (Supplementary material 3, Fig. S1). The different biology of the two web types was reflected in the taxonomic representation of taxa on sheet versus orb webs. For example, no orb webs contained nematodes or rotifers, while 20% of sheet webs contained nematodes and 65% contained rotifers. Similarly, of the nine insect orders, all were found on sheet webs, but only four on orb webs. Detailed results of OTU and taxonomic representations are accessible in Supplementary materials 3 and original data in Supplementary material 4.

**Figure 2:**
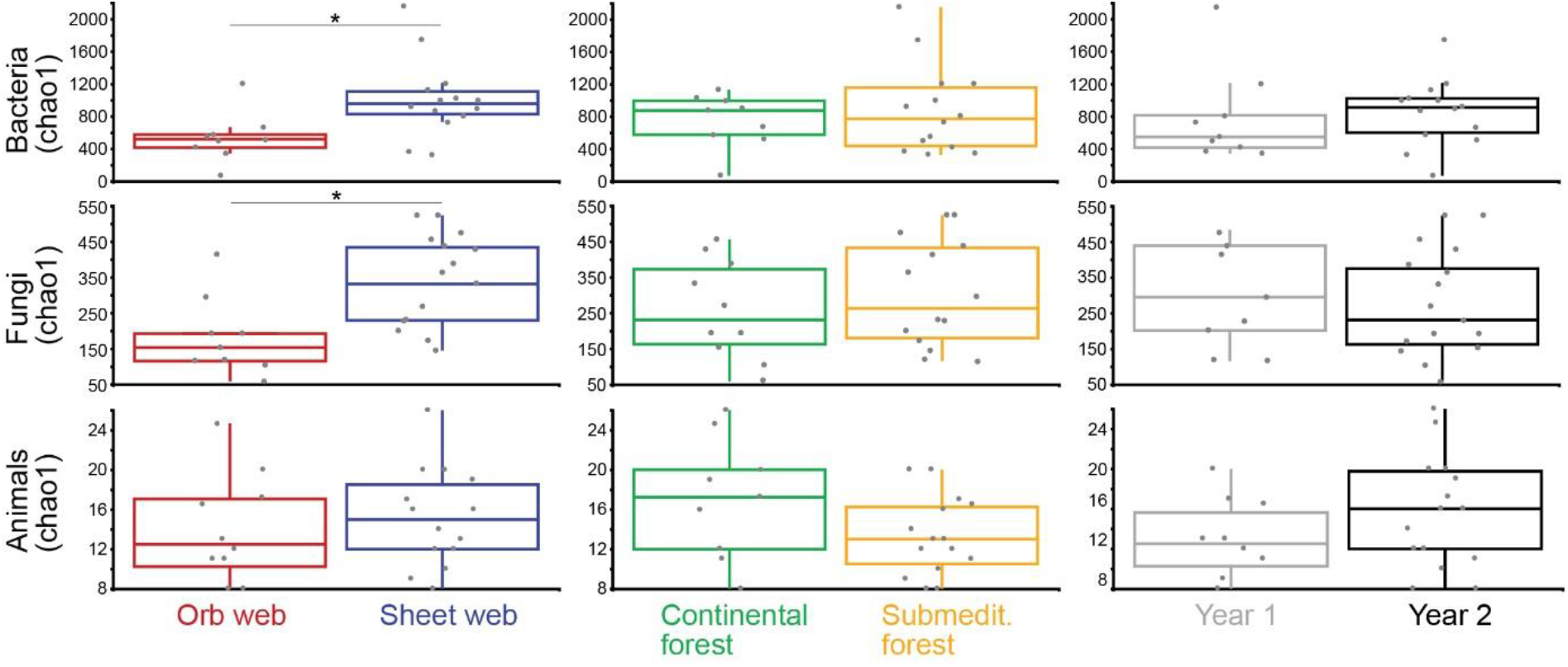
The alpha diversities of bacteria, fungi and animals, inferred from spider web eDNA using a DNA metabarcoding approach, and compared between web types (a), forests (b) and years sampled (c). Asterisks mark statistically significant differences.

Sampling locality, web type, and sampling time all affected the inferred beta diversities (community composition) of bacteria (web type: p = 0.006, *F* = 1.48165; locality: p = 0.003, *F* = 1.52083; year: p = 0.011, *F* = 1.37463), fungi (web type: p = 0.001, *F* = 5.73776; locality: p = 0.013, *F* = 2.0729; year: p = 0.001, *F* = 3.86025) and animals (web type: p = 0.008, *F* = 1.83303; locality: p = 0.001, *F* = 3.27745; year: p = 0.007, *F* = 1.7966). The PCoA plots visualize differences community compositions, and both indices using presence/absence information only (unweighted Unifrac, Jaccard index), as well as those incorporating abundance data (weighted Unifrac, Bray-Curtis coefficient), show similar results (Fig. 3; Supplementary material 3, Fig. S2).

**Figure 3:**
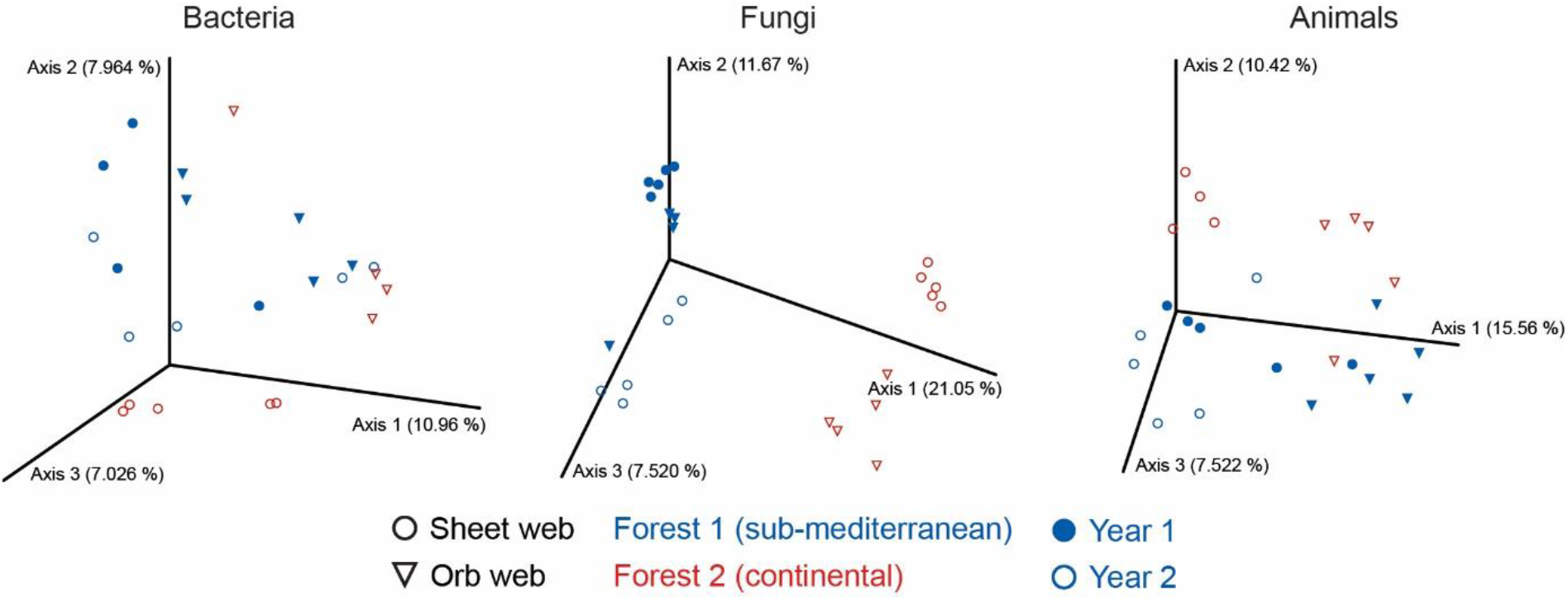
The beta diversities (community composition) of bacteria, fungi, and animals, inferred from spider web eDNA using a DNA metabarcoding approach. The PCoA plots visualize the beta diversities estimated using the unweighted Unifrac distance metric.

## Discussion

Spider webs prove suitable for detecting genetic traces of organisms in both a targeted search of a specific species and within a metabarcoding approach. We successfully trace prey introduced into spider webs and show that the targeted search for eDNA from individuals of a single species is a valid concept. In addition, we show that spider webs act as passive filters of the air column by accumulating genetic traces of diverse organisms. These results suggest spider webs are a promising tool for general biodiversity monitoring through DNA metabarcoding.

The idea of using organisms and their extended phenotypes, such as spiders and their webs, to sample genetic material of other local organisms, parallels recent studies using bloodsucking insects to identify the local fauna. For example, mosquito blood had traces of their avian, mammalian and amphibian hosts^45^, tick blood was used to identify their rodent and stoat hosts^46^, blood from leeches identified a range of mammalian species^47^, and carrion flies successfully identified diverse mammals^48^. Similarly, the contents of spider intestines can assess local arthropod biodiversity^49^. By accumulating DNA of the hosts’ prey, spider webs resemble the intestine contents of bloodsucking and predatory invertebrates. However, spiders are generalist predators, and furthermore their webs are passive traps^34^. We therefore expected that spider webs would accumulate a more general sample of organisms compared to intestine contents. Showing that spider webs efficiently capture diverse “aerial plankton” from all domains of life, our results support this expectation.

### Single-species eDNA detection

Our tests used two spider species that construct different webs. The capture threads of the hammock web are dry, while the garden spider’s classical orb contains glue-coated spirals. Although this would intuitively suggest more prey detection in the gluey web, this was not the case. The equally successful detection of prey in all our samples (Fig. 1b) rather indicates that even the smallest traces of arthropods in webs allow for successful amplification. Furthermore, our approach allows the tracing of DNA of even the smallest-bodied arthropods. Having fed spiders 5 mg crickets, we used a size class below which potential prey is often ignored due to low nutritional value^50–53^. Both these arthropod “leftovers”, as well as larger prey are thus expected to be easily detected.

Since spiders tend to cut fallen debris out of their webs, but usually ignore the presence of small, dust-like particles such as plant pollen, fungal spores, and bacteria, these organisms should also be detectable as single-species eDNA in spider webs. While each spider web in our experiments represented a single sample, future studies could pool several webs in a habitat of interest, thereby even increasing the detection power of such approach.

### DNA metabarcoding

A good eDNA tool for assessing local biodiversity should result in beta diversity reflecting differences in sample types and sample collection. Our results show that forest type (sub-mediterranean versus continental), time of collection (same season in two consecutive years), and web type (sticky and rebuilt daily versus non-sticky and long-lasting) all affect the recovered beta diversity of animal, fungal and bacterial communities. Thus, the information obtained from a few individual webs appears to be sufficient to show differences in community compositions among and even within habitats (i.e. among web types). The differentiation of samples based on web type, locality and year is particularly pronounced for fungal communities (Fig. 4). This result is in accordance with the research on fungal communities of soil samples, where metabarcoding of fungal taxa has high discrimination power^54^. Similarly, the airborne microbiomes fluctuate in time and space, and our data thus likely reflects actual differences rather than stochastic variability^55,56^. Thus, in particular, our results indicate that eDNA from spider webs could be useful in tracking biodiversity in time and space. For example, using a general sampling of spider webs, general biodiversity monitoring could be carried out throughout a season or over several years. On the other hand, by sampling spider webs in specific microhabitats, i.e. vegetation layers, useful spatial information could be brought into research of community compositions.

### Utility of eDNA from spider webs

When using eDNA to draw conclusions about the proximity of organisms relative to the traces of their DNA, it is crucial to also consider the spatial and temporal distribution and persistence of eDNA. For example, traces of genetic material remain in water for up to several weeks, but can persist for decades or even hundreds of thousands of years in soil or in permafrost^3,13,14^. The fast degradation of DNA in aquatic ecosystems makes eDNA useful in addressing topics like nature conservation, where positive detection reflects the contemporary presence of species and populations. However, in aquatic eDNA samples, the release of older genetic material from bottom sediments and the transport of eDNA in flowing and marine waters are potential sources of contamination^3,9^. Similar to aquatic ecosystems, eDNA collected from the air column via spider webs represents a contemporary presence of organisms in the environment, with potentially more accurate spatial and temporal information. Also, we find temporal contaminations unlikely. For example, some web types (e.g. sheet webs) are suspended for several weeks or even months, while most orb webs are rebuilt daily^33,34^. In addition, many spider species choose web-specific microhabitats^34^. While further studies are needed to elucidate the degeneration rate of genetic traces on longer lasting spider webs, it is clear that spider webs are not only a new tool for “filtering out” genetic material from the air column, but are also unique in the precise spatial and temporal information they provide.

In a straight forward application, eDNA from spider webs could provide a non-invasive and simple method for identifying juvenile spider specimens that cannot be determined morphologically, and could lead to new ways of studying interactions between spiders and their prey without the labor-intensive and biased sampling of potential target prey^30^. However, our results show a much broader utility of eDNA from spider webs. For example, webs could be used to address questions related to entire communities, such as the distribution and composition of arthropods, plants, fungi and bacteria over seasons and years, habitats, etc. Immediate applications in nature conservation are also foreseeable, examples being the assessment of declining global insect biodiversity^57–59^. More precisely, the development of a general pollinator eDNA sampling approach could be used as an information platform to counter the concerning, global pollinator declines exacerbated by the lack of taxonomic expertise^60^. Indeed, among the high diversity of taxa encountered in our web samples, we found several pollinator species such as bees, flies, wasps and beetles, and including the endangered longhorn beetles (Supplementary material 3). In addition to tracking pollinators and endangered species, eDNA from spider webs could also be used to track invasive or pathogen-carrying mosquitoes^61^, perhaps as an alternative to direct sampling of water puddles. Furthermore, our results indicate that eDNA from spider webs could be used to investigate species associations. For example, we found co-occurring taxa that are in known relationships, both parasitic and mutualistic (Supplementary material 3).

In our samples, we found several animal, fungal and bacterial taxa that are of agricultural and medical importance to humans (Supplementary material 3), which highlights a range of possible uses of eDNA from spider webs. Notable examples among plant pathogens and disease-causing agents are genera that include grain rust and wheat curl mites (Eriophyidae)^62,63^, gall and fungus gnats (Cecidomyiidae, Sciaridae)^64,65^, aphids (e.g. *Pineus* and *Anoecia*)^66–68^, several fungal representatives that damage wheat crops (e.g. *Fusarium* and *Ustilago*)^69–71^, eudicot plants (*Verticillium*)^72^, and rice (*Magnaporthe grisea*)^73^, as well as several bacterial genera that include plant pathogenic representatives, e.g. *Pseudomonas*, *Erwinia*, *Dickea*, and *Pectobacterium*^74^. Therefore, spider web eDNA could be used for early detection of agricultural pests, even in the absence of disease symptoms, an approach already demonstrated for eDNA from orchards^75–77^. Moreover, we found several medically important fungal and bacterial representatives, many of which cause respiratory problems and allergies (e.g. *Aspergillus*, *Stachybotrys* and *Botrytis*)^78–80^, food spoilage and gastrointestinal infections (e.g. *Absidia*)^81^, or are associated with livestock (*Salmonella* and *Clostridium*)^82^ and invertebrate vectors (e.g. *Haemophilus*)^83,84^. Perhaps, future use of eDNA from spider webs can extend our understanding about the presence and role of human pathogens in non-urban environments.

## Concluding remarks

Spider webs are passive air filters. Utilizing them as an eDNA source thus represents a novel technique for sampling air masses, an approach equivalent to sampling other media such as water, ice, and sediments. Our results show that spider webs can become a new tool for the targeted tracking and general monitoring of any kind of organisms, from arthropods, rotifers and fungi, to bacteria, plants and perhaps viruses. As such, the use of eDNA from spider webs offers numerous potential applications from biodiversity monitoring, tracking invasive and pest species, animal diet assessment, obtaining climate change data, to studies on the distribution and niches of arthropods, plants, fungi and bacteria, whether in a single species or in a metabarcoding context.

## Supplementary material

Supplementary material 1: Additional literature overview.

Supplementary material 2: Additional methods: DNA isolation protocol, primer details and amplification protocols.

Supplementary material 3: Additional metabarcoding results: alpha and beta diversity results, and comments on taxa of human interest.

Supplementary material 4: QIIME2 output files (view at https://view.qiime2.org/ or unzip for raw data).

## Acknowledgements

We thank the technical and student staff at the Department of Biotechnology and Systems Biology, National Institute of Biology, for their logistic and technical help in the laboratory. We thank S. Koren and Omega d.o.o. for their help with the microfluidic capillary electrophoresis. This study was funded by the following grants from the Slovenian Research Agency: Z1-8143, P1-0236, P4-0407, P1-0198, P1-0255, J1-9163 and J1-1703.

## Author contributions

M.G., M.K., D.K., and M.R. conceived the study. M.G., K.B., A.P., and C.G. performed experiments and analyses. All authors jointly wrote the paper.

## Data availability

All experimental data are available in the manuscript and supplementary material 4.

## Supplementary material 1: Additional literature overview

## 1. Additional literature overview

Recently, spider webs have been proposed as a novel source of eDNA. Xu et al.^1^ maintained two black widow species (*Latrodectus hesperus* and *L. mactans*) in controlled laboratory conditions, feeding them a single prey type, the house cricket *Acheta domestica*. Using conventional PCR, they detected genetic traces of host and prey, and both spider and prey DNA remained detectable for months^1^. Blake et al.^2^ confirmed these findings by amplifying spider DNA from silk of the laboratory kept bird spider *Psalmopoeus*, and from the daddy long leg spider *Pholcus phalangioides*, sampled in a house. Additionally, in a study investigating ways of improving the taxonomic coverage in DNA metabarcoding, Corse et al.^3^ included spider webs as one of several sources of eDNA. Because spider webs are ubiquitous and can be easily collected, the potential use of web derived eDNA goes well beyond identifying spiders and their prey. For example, spider webs are known to accumulate plant pollen and agrochemical spray^4,5^. Thus, utilizing the genetic material obtained from spider webs represents an alternative, novel, and potentially powerful sampling method to supplement and complement traditional methods.

While these studies are valuable for introducing the concept of eDNA from spider webs, they urgently call for a focused investigation of the utility of spider webs as an eDNA source. Specifically, the two studies^1,2^ targeting host and prey DNA in a single-species approach, suffer from two major shortcomings. First, sampling webs in controlled conditions means that factors like UV light, heat, humidity, wind, and rain have likely been reduced compared to field conditions. Second, these two studies failed to apply standard laboratory controls, e.g. controlling for possible isolation inhibition, false positive/negative results, and optimal assay performance throughout a wide concentration of isolated target DNA^6–8^. Furthermore, Corse et al.^3^ show that spider webs accumulate a diversity of arthropod genetic traces, but their study focused on identification resolution of different metabarcoding approaches for eDNA from different environments, rather than spider webs as an eDNA source, *per se*. Here, we first tested the detection of a model arthropod target in the field. Second, using a metabarcoding approach, we investigated the utility of spider web eDNA in general biodiversity monitoring.

## Supplementary material 2: Additional methods

## 1. Protocol for DNA isolation

This protocol is an adapted protocol of the PowerLyzer^®^ PowerSoil^®^ DNA Isolation Kit (Qiagen, USA).

1. Add sample into provided PowerBead tube.
2. Add 750 μl of bead solution.
3. Vortex briefly, and briefly spin-down centrifuge.
4. Add 60 μl of C1 solution. If C1 solution precipitated, heat to 60°C until dissolved.
5. Vortex briefly or invert tube several times.
6. Homogenize using FastPrep (XX): 45 s, MP adapter (24x), 6.5 ms.
7. Incubate at 70°C for 10 min while shaking at 2000 rpm.
8. Homogenize using FastPrep (XX): 45 s, MP adapter (24x), 6.5 ms.
9. Centrifuge at 10,000 x g, 2 x for 1 min at room temperature, in between tap with fingers to remove foam.
10. Transfer 400-500 μl of supernatant into new 2 ml collection tube. Avoid transferring parts of pellet. If there is not enough supernatant, repeat centrifuge at 10,000x g for 1 min at room temperature.
11. Add 250 μl of C2 solution to supernatant.
12. Vortex briefly.
13. Incubate at 4°C for 5 min.
14. Centrifuge at 10,000 x g for 1 min at room temperature.
15. Transfer 600 μL of supernatant into new 2 ml collection tube. Avoid transferring parts of pellet.
16. Add 200 μl of solution C3.
17. Vortex briefly.
18. Incubate at 4°C for 5 min.
19. Centrifuge at 10,000 x g for 1 min at room temperature.
20. Transfer 625μl of supernatant into new 2 ml collection tube. Avoid transferring parts of pellet.
21. Add 1000 μl of solution C4 to the supernatant.
22. Vortex for 5 s, spin-down centrifuge briefly.
23. Transfer 600 μl of supernatant to a spin filter.
24. Centrifuge at 10,000 x g for 1 min at room temperature.
25. Discard the flow-through and place spin filter into clean 2 ml collection tube.
26. Add an additional 600 μl to spin filter.
27. Centrifuge at 10,000 x g for 1 min at room temperature.
28. Discard the flow-through and place spin filter into clean 2 ml collection tube.
29. Add the remaining supernatant on spin filter.
30. Centrifuge at 10,000 x g for 1 min at room temperature.
31. Discard the flow-through and place spin filter into clean 2 ml collection tube.
32. Add 500 μl of solution C5.
33. Centrifuge at 10,000 x g for 30 s at room temperature.
34. Discard the flow-through and place spin filter into clean 2 ml collection tube.
35. Centrifuge at 10,000 x g for 1 min at room temperature.
36. Discard the flow-through and place spin filter into clean 2 ml collection tube.
37. Add 75 μl of solution C6 to the center of the filter membrane, without touching membrane.
38. Incubate at room temperature for 3-5 min.
39. Centrifuge at 10,000 x g for 30 s at room temperature.
40. Discard the spin filter, keep DNA frozen.

## 2. Single-species eDNA detection

**Table S1:**
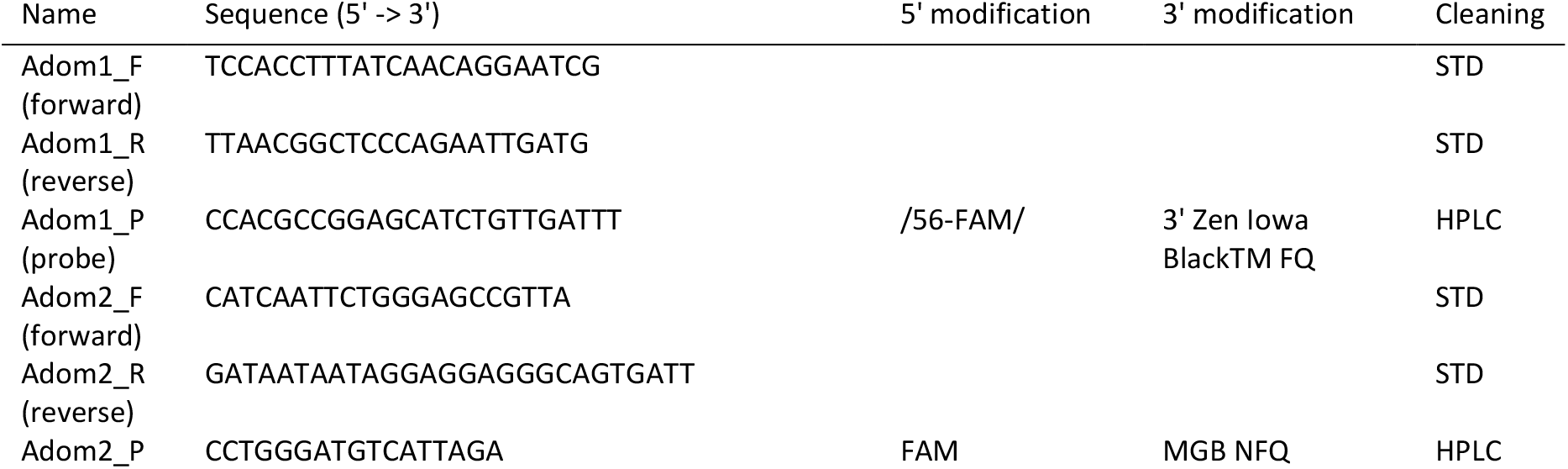
Adom1 and Adom2 primer details.

**Table S2:** Reaction mixes for qPCRs using Adom1 primers. We added 2 μl of isolated DNA sample for a 10 μl total qPCR.

| Adom1 mastermix | Volume per reaction (µl) |
| --- | --- |
| Water | 0.2 |
| 2× TaqMan Universal master mix (ABI) | 5 |
| Adom1_F (10 µM) | 0.9 |
| Adom1_R (10 µM) | 0.9 |
| Adom1_P (2.5 µM) | 1 |
| Sum | 8.0 |

**Table S3:**
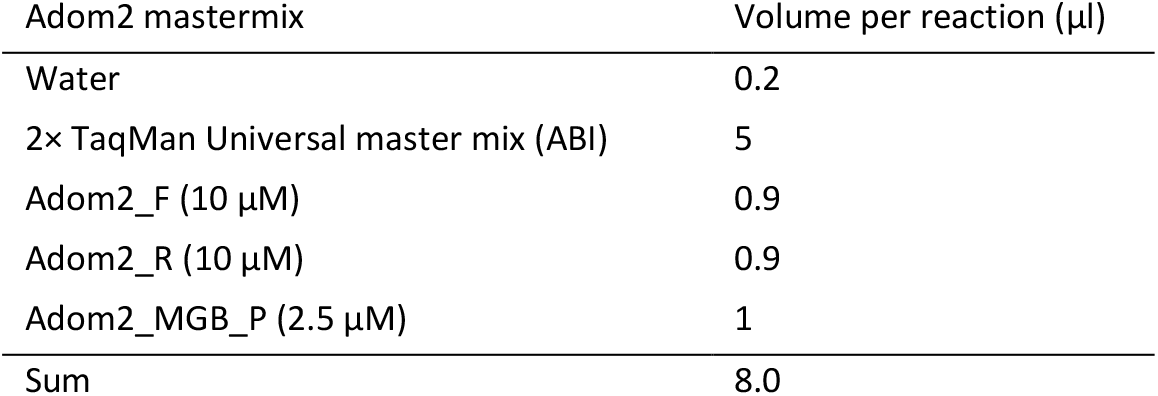
Reaction mixes for qPCRs using Adom2 primers. We added 2 μl of isolated DNA sample for a 10 μl total qPCR.

**Table S4:** Reaction mixes for qPCRs using the “Eukaryotic 18S rRNA Endogenous Control” (Applied Biosystems, USA). We added 2 μl of isolated DNA sample for a 10 μl total qPCR.

| Adom2 mastermix | Volume per reaction (µl) |
| --- | --- |
| Water | 2.5 |
| 2× TaqMan Universal master mix (ABI) | 5 |
| Primers/probes (20 x) | 0.5 |
| Sum | 8.0 |

**Table S5:**
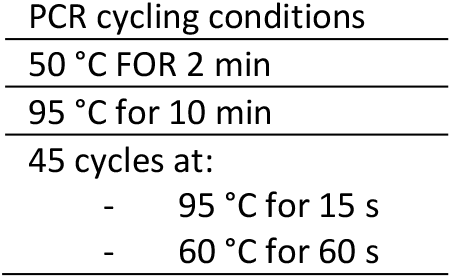
qPCR amplification protocol for Adom1 amplification, using the 7900HT Fast Real-Time PCR System (Applied Biosystems, USA).

|  |
| --- |
| PCR cycling conditions |
| 50 °C FOR 2 min |
| 95 °C for 10 min |
| 45 cycles at: |
| - 95 °C for 15 s |
| - 60 °C for 60 s |

## 3. DNA metabarcoding

**Table S6:** Sampling details for the metabarcoding experiment. The sample codes (A, B, C) for each sample correspond to Qiime2 visual output files (Supplementary material 3).

| Animals | Fungi | Bacteria | Web type | Forest type | Locality | Coordinates | Sampling date |
| --- | --- | --- | --- | --- | --- | --- | --- |
| A01 | B01 | C01 | Sheet web | Submediterranean | Locality 1 | 45.575873,<br>13.856277 | 5.x.2018 |
| A02 | B02 | C02 | Sheet web | Submediterranean | Locality 1 | 45.575873,<br>13.856277 | 5.x.2018 |
| A03 | B03 | C03 | Sheet web | Submediterranean | Locality 1 | 45.575873,<br>13.856277 | 5.x.2018 |
| A04 | B04 | C04 | Sheet web | Submediterranean | Locality 1 | 45.575873,<br>13.856277 | 5.x.2018 |
| A05 | B05 | C05 | Sheet web | Submediterranean | Locality 1 | 45.575873,<br>13.856277 | 5.x.2018 |
| A06 | B06 | C06 | Sheet web | Continental | Locality 2 | 46.515795,<br>15.690766 | 28.ix.2018 |
| A07 | B07 | C07 | Sheet web | Continental | Locality 2 | 46.515795,<br>15.690766 | 28.ix.2018 |
| A08 | B08 | C08 | Sheet web | Continental | Locality 2 | 46.515795,<br>15.690766 | 28.ix.2018 |
| A09 | B09 | C09 | Sheet web | Continental | Locality 2 | 46.515795,<br>15.690766 | 28.ix.2018 |
| A10 | B10 | C10 | Sheet web | Continental | Locality 2 | 46.515795,<br>15.690766 | 28.ix.2018 |
| A11 | B11 | C11 | Orb web | Continental | Locality 2 | 46.515795,<br>15.690766 | 28.ix.2018 |
| A12 | B12 | C12 | Orb web | Continental | Locality 2 | 46.515795,<br>15.690766 | 28.ix.2018 |
| A13 | B13 | C13 | Orb web | Continental | Locality 2 | 46.515795,<br>15.690766 | 28.ix.2018 |
| A14 | B14 | C14 | Orb web | Continental | Locality 2 | 46.515795,<br>15.690766 | 28.ix.2018 |
| A15 | B15 | C15 | Orb web | Continental | Locality 2 | 46.515795,<br>15.690766 | 28.ix.2018 |
| A16 | B16 | C16 | Sheet web | Submediterranean | Locality 1 | 45.575873,<br>13.856277 | 26.ix.2017 |
| A17 | B17 | C17 | Sheet web | Submediterranean | Locality 1 | 45.575873,<br>13.856277 | 26.ix.2017 |
| A18 | B18 | C18 | Sheet web | Submediterranean | Locality 1 | 45.575873,<br>13.856277 | 26.ix.2017 |
| A19 | B19 | C19 | Sheet web | Submediterranean | Locality 1 | 45.575873,<br>13.856277 | 26.ix.2017 |
| A20 | B20 | C20 | Sheet web | Submediterranean | Locality 1 | 45.575873,<br>13.856277 | 26.ix.2017 |
| A21 | B21 | C21 | Orb web | Submediterranean | Locality 1 | 45.575873,<br>13.856277 | 26.ix.2017 |
| A22 | B22 | C22 | Orb web | Submediterranean | Locality 1 | 45.575873,<br>13.856277 | 26.ix.2017 |
| A23 | B23 | C23 | Orb web | Submediterranean | Locality 1 | 45.575873,<br>13.856277 | 26.ix.2017 |
| A24 | B24 | C24 | Orb web | Submediterranean | Locality 1 | 45.575873,<br>13.856277 | 26.ix.2017 |
| A25 | B25 | C25 | Orb web | Submediterranean | Locality 1 | 45.575873,<br>13.856277 | 26.ix.2017 |

**Table S7:** The number of obtained reads for all samples. Red lines mark the omitted samples, i.e. the chosen sample/amplicon combination for beta diversity estimation.

| Sample ID | Animal reads | Sample ID | Fungi reads | Sample ID | Bacteria reads |
| --- | --- | --- | --- | --- | --- |
| A12 | 31797 | B01 | 94101 | C06 | 140830 |
| A11 | 23215 | B09 | 66780 | C09 | 128351 |
| A13 | 22053 | B04 | 63497 | C13 | 128288 |
| A24 | 20934 | MockComm | 56655 | C23 | 125484 |
| A14 | 14717 | B18 | 56564 | C01 | 123588 |
| A01 | 14345 | B19 | 53879 | MockComm | 123466 |
| A23 | 7330 | B07 | 46441 | C20 | 122900 |
| A05 | 7137 | B10 | 46312 | C14 | 121516 |
| A21 | 5262 | B02 | 43853 | C03 | 119455 |
| A17 | 4930 | B06 | 40036 | C10 | 117504 |
| A15 | 3431 | B08 | 38681 | C07 | 115460 |
| A08 | 2745 | B25 | 35287 | C18 | 111457 |
| A22 | 2710 | B03 | 32613 | C17 | 107573 |
| A18 | 2162 | B05 | 15780 | C04 | 106685 |
| A25 | 1704 | B13 | 14283 | C25 | 99389 |
| A16 | 1654 | B17 | 13272 | C05 | 96528 |
| A02 | 1648 | B21 | 13196 | C08 | 96393 |
| A04 | 1281 | B15 | 11228 | C02 | 95965 |
| A10 | 1021 | B20 | 11182 | C21 | 81834 |
| A19 | 978 | B16 | 8410 | C24 | 35137 |
| A06 | 971 | B12 | 7862 | C19 | 32071 |
| A03 | 790 | B23 | 3160 | C12 | 25121 |
| A07 | 584 | B24 | 1976 | C22 | 16941 |
| A20 | 491 | B14 | 1840 | C15 | 15906 |
| A09 | 116 | B11 | 1361 | Neg-4 | 66 |
| MockComm | 0 | B22 | 0 | Neg-1 | 8 |
| Neg-1 | 0 | Neg-1 | 0 | Neg-3 | 6 |
| Neg-2 | 0 | Neg-2 | 0 | C11 | 3 |
| Neg-3 | 0 | Neg-3 | 0 | C16 | 1 |
| Neg-4 | 0 | Neg-4 | 0 | Neg-2 | 1 |

**Table S8:**
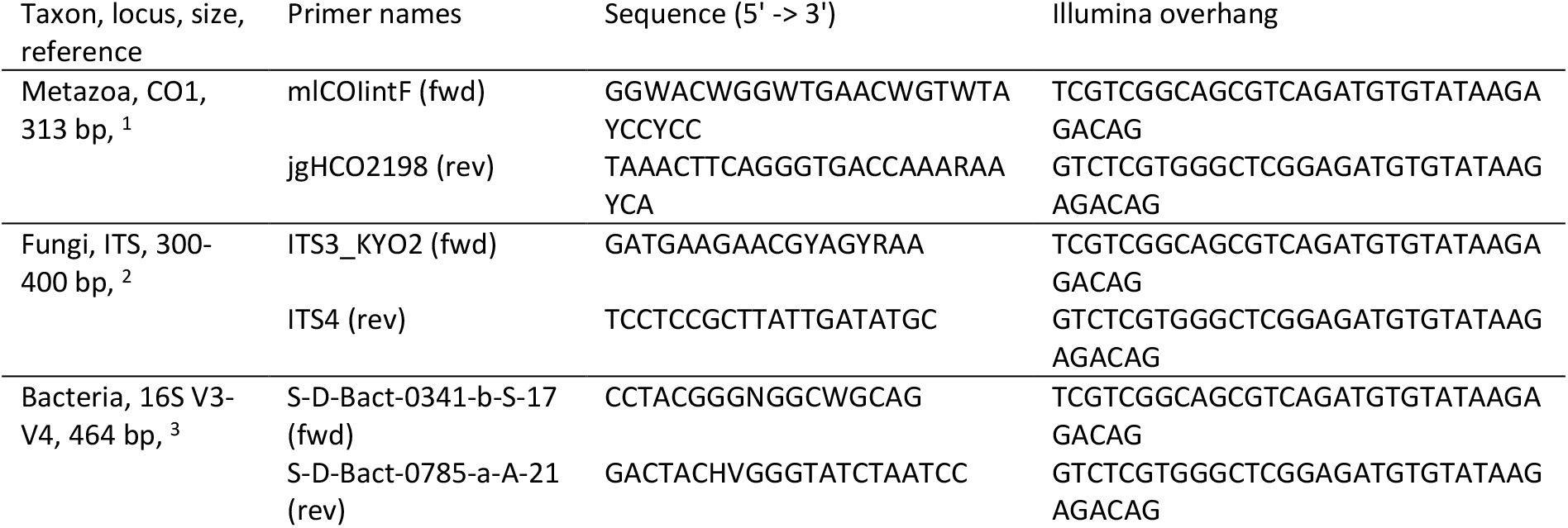
Primers used in DNA metabarcoding.

**Table S9:** PCR protocol for amplification of metabarcoding primers. To avoid amplification biases, we chose an amount of cycles where no reactions were saturated. We estimated PCR saturation using gel electrophoresis, i.e. we chose conditions where DNA bands on the gel were not or barely visible.

|  |
| --- |
| PCR cycling conditions |
| 95 °C for 3 min |
| 20 cycles at: |
| - 95 °C for 30 s |
| - 55 °C for 30 s |
| - 72 °C for 30 s |
| 72 °C for 5 min |
| Hold at 4 °C |

## Supplementary material 3: Additional DNA metabarcoding results

## 1. Alpha diversity: additional results

The statistical analysis of the Shannon index of alpha diversity showed that in the two forests, webs accumulate an equal diversity of bacterial (p = 0.208, H = 1.587), fungal (p = 0.198, H = 1.659) and animal (p = 0.325, H = 0.968) eDNA. The sampling year yielded a similar diversity of eDNA (bacteria: p = 0.413, H = 0.671; fungi: p = 0.200, H = 1.644; animals: p = 0.292, H = 1.111). Compared to orb webs, sheet webs accumulated a higher diversity of bacterial (p = 0.051, H = 3.813) and fungal (p = 0.025, H = 5.000) eDNA, but not animal eDNA (p = 0.089, H = 2.883).

## 2. Number of OTUs and lists of taxa

**Figure 1:**
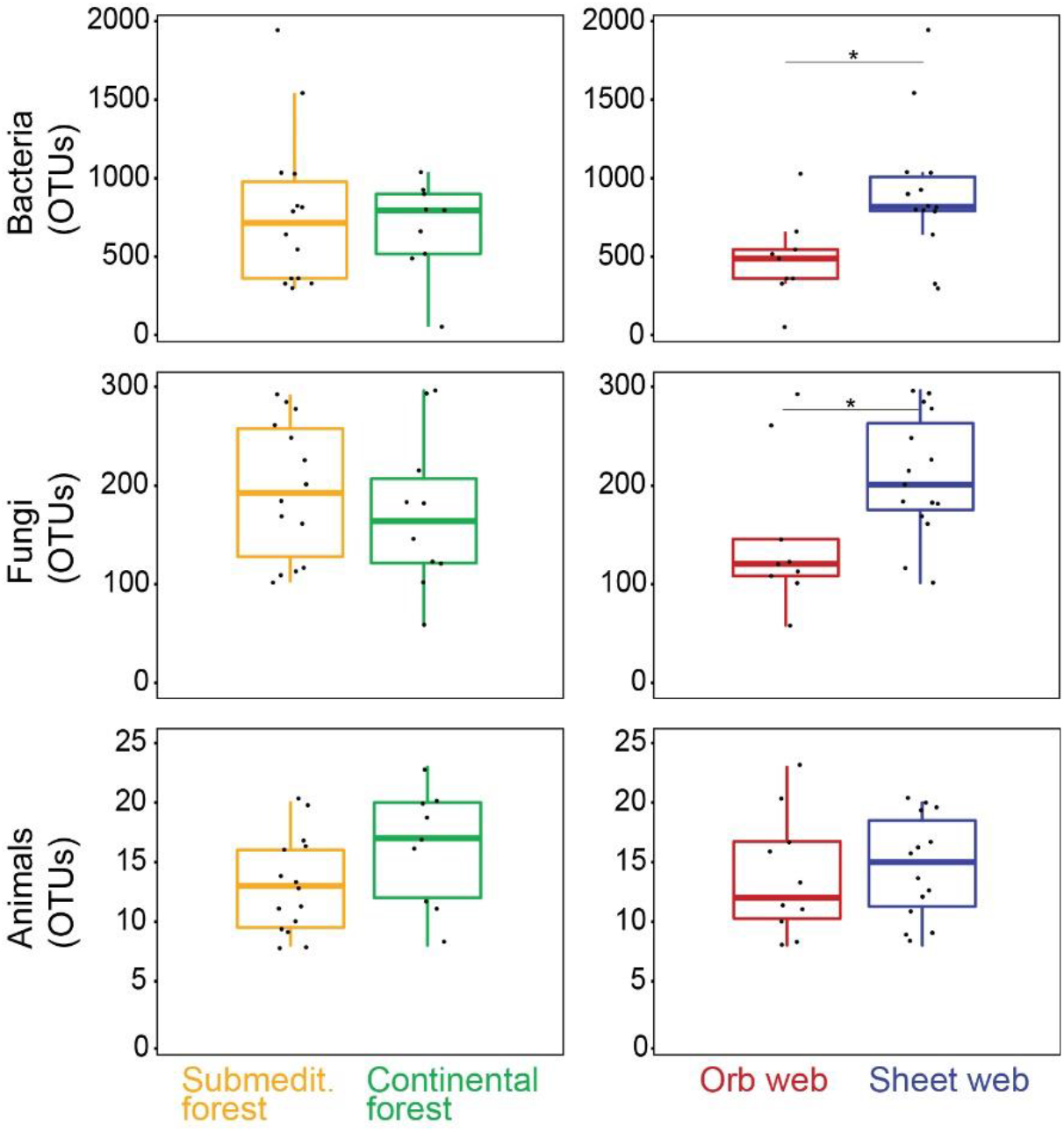
Average operational taxonomic unit (OTU) number per web. Data are shown for three different taxonomic groups (Bacteria, Fungi, Animals) and grouped by factors designated by color. The distribution of values for each factor is represented by box-whisker plots. In the two forests, webs on average accumulated an equal number of bacterial (p = 0.928, U = 61), fungal (p = 0.596, U = 60.5) and animal (p = 0.128, U = 41.5) OTUs. Compared to orb webs, sheet webs on average accumulated more bacterial (p = 0.018, U = 25) and fungal (p = 0.039, U = 32.5), but not animal (p = 0.598, U = 60.5) OTUs. Asterisks mark statistically significant differences.

**Table 1:** List of identified bacterial phyla with OTU counts, cumulative for all samples.

| Phylum | OTUs | Phylum | OTUs |
| --- | --- | --- | --- |
| Acidobacteria | 424 | Fusobacteria | 8 |
| Actinobacteria | 1184 | Gemmatimonadetes | 148 |
| Armatimonadetes | 247 | Hydrogenedentes | 1 |
| Bacteroidetes | 1699 | Kiritimatiellaeota | 1 |
| BRC1 | 23 | Lentisphaerae | 2 |
| Chlamydiae | 51 | Nitrospirae | 4 |
| Chloroflexi | 297 | Patescibacteria | 134 |
| Cyanobacteria | 156 | Planctomycetes | 965 |
| Deferribacteres | 2 | Proteobacteria | 2973 |
| Deinococcus-Thermus | 29 | Rokubacteria | 1 |
| Dependentiae | 10 | RsaHF231 | 1 |
| Elusimicrobia | 4 | Tenericutes | 9 |
| Entotheonellaeota | 2 | Verrucomicrobia | 604 |
| Epsilonbacteraeota | 2 | WPS-2 | 15 |
| FBP | 87 | N/a | 1710 |
| Firmicutes | 492 | Total | 11285 |

**Table 2:** A list of identified fungal phyla, cumulative for all samples, with OTU counts, and lists of all identified classes of Ascomycota and Basidiomycota, with OTU counts, respectively.

| Fungi |  | Ascomycota |  | Basidiomycota |  |
| --- | --- | --- | --- | --- | --- |
| Phylum | OTUs | Class | OTUs | Class | OTUs |
| Ascomycota | 2392 | Arthoniomycetes | 12 | Agaricomycetes | 960 |
| Basidiomycota | 1440 | Dothideomycetes | 962 | Agaricostilbomycetes | 13 |
| Blastocladiomycota | 1 | Eurotiomycetes | 258 | Atractiellomycetes | 3 |
| Chytridiomycota | 15 | Lecanoromycetes | 107 | Cystobasidiomycetes | 51 |
| Entomophthoromycota | 2 | Leotiomycetes | 280 | Dacrymycetes | 3 |
| Mortierellomycota | 7 | Orbiliomycetes | 74 | Exobasidiomycetes | 50 |
| Mucoromycota | 10 | Pezizomycetes | 23 | Malasseziomycetes | 2 |
| Rozellomycota | 1 | Saccharomycetes | 33 | Microbotryomycetes | 44 |
| Zoopagomycota | 1 | Sordariomycetes | 412 | Pucciniomycetes | 31 |
| N/a | 136 | Taphrinomycetes | 12 | Spiculogloeomycetes | 5 |
| Total | 4005 | N/a | 219 | Tremellomycetes | 193 |
|  |  | Total | 2392 | Ustilaginomycetes | 14 |
|  |  |  |  | Wallemiomycetes | 8 |
|  |  |  |  | N/a | 63 |
|  |  |  |  | Total | 1440 |

**Table 3:** Number of classes, orders, and families within each fungal phylum, cumulative for all samples.

| Phylum | Classes | Orders | Families |
| --- | --- | --- | --- |
| Ascomycota | 10 | 57 | 167 |
| Basidiomycota | 13 | 43 | 130 |
| Blastocladiomycota | 1 | 1 | 1 |
| Chytridiomycota | 1 | 1 | 1 |
| Entomophthoromycota | 1 | 1 | 1 |
| Mortierellomycota | 1 | 1 | 1 |
| Mucoromycota | 2 | 2 | 4 |
| Rozellomycota | 1 | 1 | 1 |
| Zoopagomycota | 1 | 1 | 1 |
| Total | 31 | 108 | 307 |

**Table 4:** Lists of identified animal phyla, arthropod classes, and insect orders, with OTU counts, respectively. Cumulative for all samples.

| Animalia |  | Arthropoda |  | Insecta |  |
| --- | --- | --- | --- | --- | --- |
| Phylum | OTUs | Class | OTUs | Order | OTUs |
| Arthropoda | 263 | Arachnida | 37 | Coleoptera | 3 |
| Nematoda | 3 | Entognatha | 3 | Dermaptera | 1 |
| Rotifera | 48 | Insecta | 218 | Diptera | 116 |
| Total | 314 | N/a | 5 | Hemiptera | 28 |
|  |  | Total | 263 | Hymenoptera | 59 |
|  |  |  |  | Lepidoptera | 3 |
|  |  |  |  | Thysanoptera | 2 |
|  |  |  |  | N/a | 6 |
|  |  |  |  | Total | 218 |

**Table 5:**
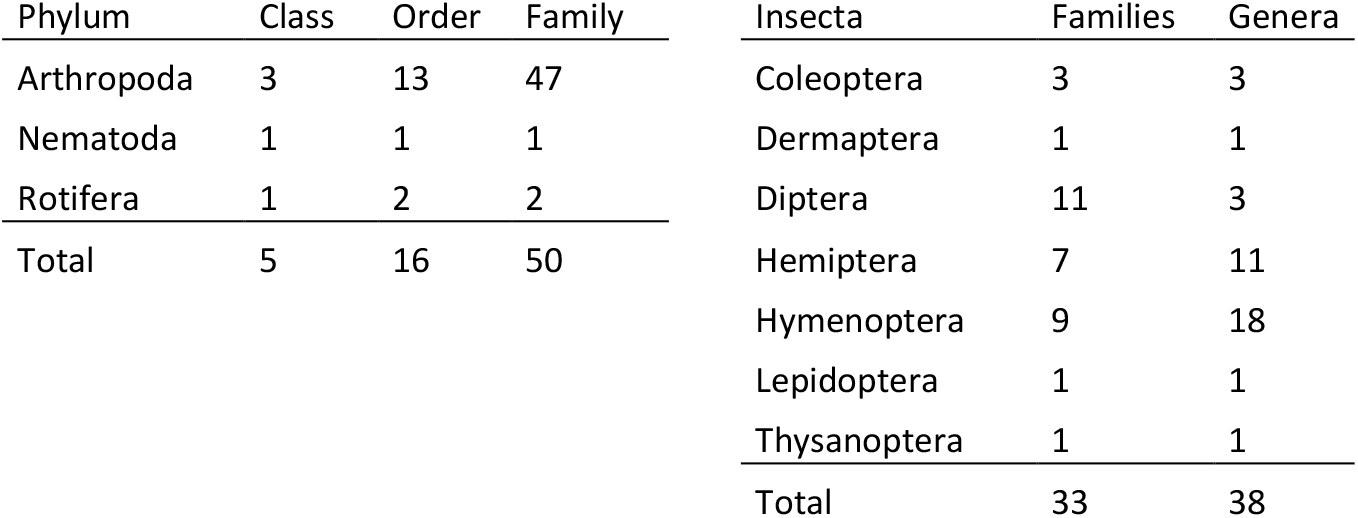
Number of classes, orders and families in animal phyla, and number of families and genera in insect orders. Cumulative for all samples.

## 3. Beta diversity: additional results

**Figure 2:**
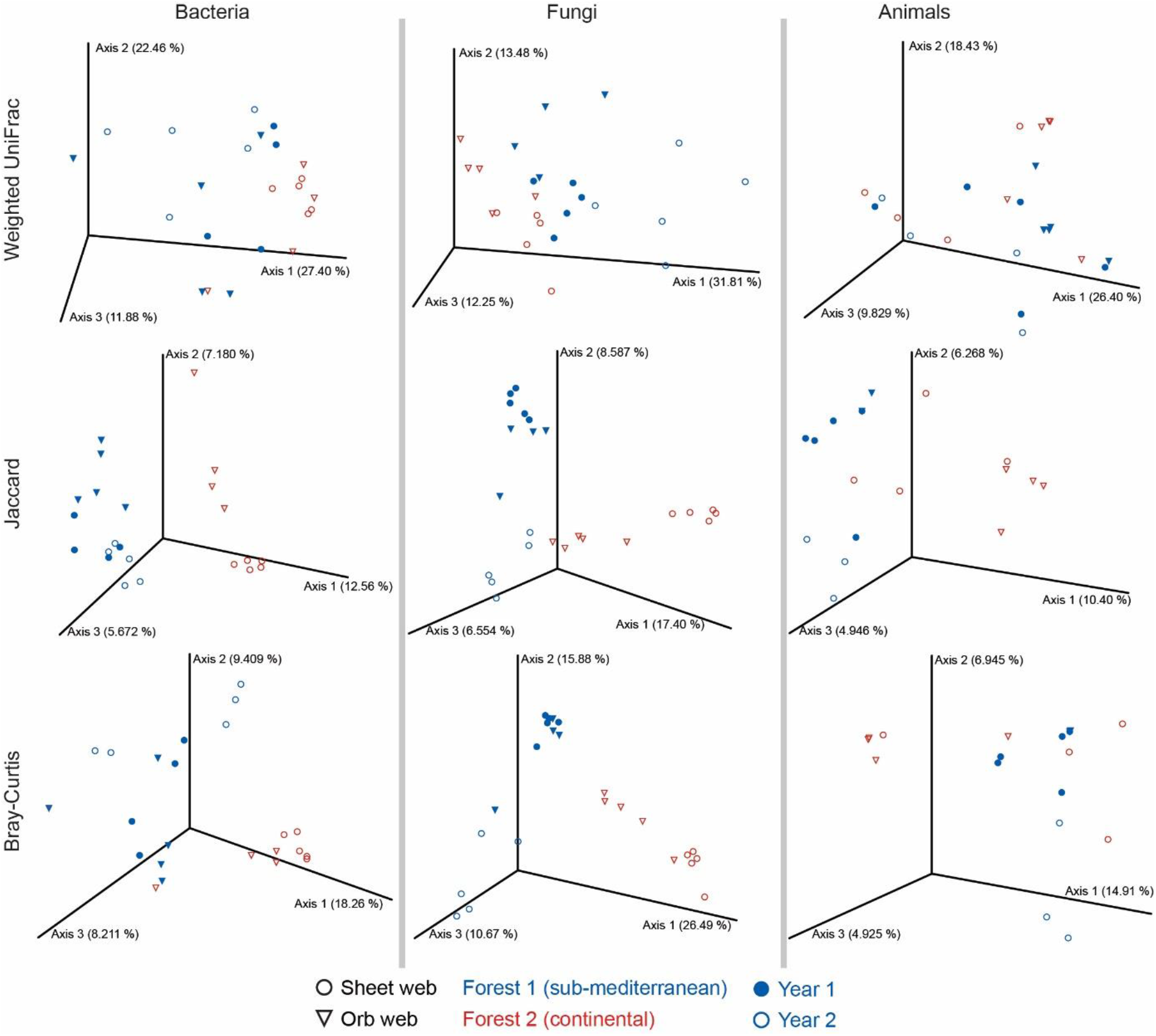
The beta diversities (community composition) of bacteria, fungi, and animals, inferred from spider web eDNA using a DNA metabarcoding approach. The PCoA plots visualize the beta diversities estimated using the weighted UniFrac distance metric, Jaccard index and the Bray-Curtis coefficient.

## 4. Animal taxa of interest

Among animals, taxa of agricultural interest are perhaps the most common. We found several representatives of gall mites, family Eriophyidae (Trombidiformes). These are plant parasites, commonly causing galls and other damage to plant tissues^1^. Specifically, we found mites of the genus *Abacarus*, which includes the grain rust mite *A. hystrix* that lives on grass and can cause big loses in crop yields^2^, and the wheat curl mite *Aceria tosichella* that is a global cereal pest and a vector of viruses like wheat streak mosaic virus^3^. Within the family Phytoptidae (Trombidiformes), we found *Nalepella brewerianae* that is a parasite of *Picea* trees, and *Phytoptus* mites that induce galls on hazel trees^4^.

Among insects, all orders include agricultural pests and it is not surprising we found several in our samples. Among Diptera, we found the leaf miner fly *Phytomyza crassiseta* (family Agromyzidae)^5^, *Delia* flies (family Anthomyiidae) whose larvae cause agricultural losses by tunneling into roots and stems of host plants^6^, and several representatives of gall gnats (Cecidomyiidae) and fungus gnats (Sciaridae). Many species of gall gnats are important pests^7^, and recent studies even hint at a largely underestimated diversity of this family that could be one of the most species rich animal families in general^8^. Fungus gnats are mostly mushroom pests^9^.

Among Hemiptera, we found pine aphids (*Pineus*, Agelidae) that infest several pine species and can cause large economic damage^10^, whiteflies (*Aleyrodes*, Aleyrodidae) that are a major agriculture threat in greenhouses^11^, *Aphrophora* spittlebugs that can cause damage on trees with soft wood^12^, the *Unaspis* snow scales that are pests of spindle trees^11^, phylloxera (Phylloxeridae) that are pests of grapevines^11^, and aphids (Aphidae), e.g. the *Anoecia* aphids that are wheat and corn pests^13^, and the waxy grey pine needle aphid *Schizolachnus pineti* that infests pine plantations^14^. Some encountered taxa are plant disease vectors, e.g. many aphids are known virus vectors^15^, while some others, e.g. the privet leafhopper *Fieberiella florii*, transmit bacteria^16^.

Among other insect orders, the earwig *Forficula auricularia* (Forficulidae, Dermaptera) can cause significant damage to crops and fruit orchards^17^. We also found thrips (Thripidae, Thysanoptera), many of which are important agricultural pests^17^.

We found several taxa that are pollinators, e.g. the chironomid flies, the orange muscid fly *Phaonia pallida*, soldier flies (Stratiomyidae), several honeybee species (*Apis*, Apidae, Hymenoptera), the common wasp *Vespula vulgaris* (Vespidae, Hymenoptera), and butterflies and moths (Lepidoptera)^18^. We found one representative of long horn beetles (Cerambycidae, Coleoptera), where several endangered species belong to. Flies from the family Chironomidae were common in our samples. These occur in high numbers and play an important role in aquatic ecosystems. They are also important indicator organisms used for assessing water quality^19^.

We found several other interesting animal representatives. For example, most sheet web samples contained traces of freshwater rotifers from the class Bdelloidea. These microscopic animals live in different water bodies, in moist soil, and on mosses and lichens on tree trunks. During dry and harsh conditions, these rotifers enter a state of dormancy^20,21^. We hypothesize that rotifers in this state could have easily been carried around by wind and picked up by spider webs. We also found rhabditid nematodes (Nematoda, Rhabditida), a group consisting of mostly zooparasitic and phytoparasitic representatives^22^ that could have been transmitted to spider webs via their insect and plant hosts. Among Hexapoda, many web samples contained slender springtails (Collembola, Entomobryidae). These springtails are common in many habitats, including the canopy and other vegetation^23^. We identified no springtails exclusive to soil.

Some taxa occurring in our samples indicate that eDNA from spider webs could be used to investigate species associations. For example, *Scutacarus* mites are associated with *Lasius flavus* ants^24^, and *Lasius* ants commonly attend to nymphs of *Anoecia* aphids to obtain aphid honeydew^25^. All these taxa were present in our samples. Furthermore, using eDNA from spider webs, one could perhaps track parasitoid-host interactions. For example, we found Eulophidae, Ichneumonidae, Mymaridae, Platygastridae and Trichogrammatidae that are large families of parasitoid wasps, whose hosts comprise of a large range of arthropods^11^. Among these wasps, *Polyaulon* wasps are ant mimics and parasitoids^26^, and co-occurred with several ant species in our samples. Also, big-headed flies (Pipunculidae) almost exclusively parasitize leafhoppers^27^. Within a single web sample, we found the big-headed fly *Clistoabdominalis* and its potential cicadid hosts. Another example is the ant *Solenopsis fugax* that obtains food by klepto-parasitizing larger ant colonies^28^, and we found it co-occurring with four other ant species.

## 5. Fungal taxa of interest

Several encountered families include representatives that are plant pathogens and thus important disease-causing agents of a large diversity of agricultural plants. Notable encountered examples include *Fusarium*, *Gaeumannomyces*, and *Ustilago* that damage barley and wheat crops^29–31^, *Verticillium* that damages hundreds of eudicot plants like cotton, tomatoes, potatoes, peppers, and eggplants^32^, the rice blast fungus *Magnaporthe grisea* that causes a serious rice disease^33^, and *Armillaria* that causes “white rot” root disease, mostly in trees and shrubs^34^.

Some encountered fungi are of potential human medical importance. We found several *Cladosporum* species that were present in most samples, and are one of the most common fungi isolated from air. Some species of this genus are human allergens that cause respiratory problems^35^. Other taxa of potential medical importance were less common. Among these, *Candida* can cause infections in immunocompromised individuals^36^. *Aspergillus* representative can cause allergies and are important crop pests, whose mycotoxins can cause human diseases^37^. *Cryptococcus neoformans* sometimes causes meningo-encephalitis in immunocompromised individuals^38^. The black molds *Stachybotrys* are linked to damp environments and can cause respiratory problems and headaches^39^. *Mucorales* (phylum Mucoromycota) fungi can cause mucormycosis, a group of serious infections of the face, nose and mouth^40^. The gray fungus *Botrytis* can cause asthma, hay fever and keratomycosis^41^. *Absidia* are common soil fungi, and some cause food spoilage and pulmonary and gastrointestinal infections in immunocompromised individuals^42^. *Bipolaris* causes sinusitis, asthma, and hay fever^43^. *Stachybotrys* and *Curvularia* occur in forests, cause type I allergies (asthma, hay fever, sinusitis)^44^, while the latter can also cause pneumonia, corneal and nail infections^39^. *Fusarium* causes hay fever, asthma and keratitis^45^. *Penicillium* is common on moldy food and its ingestion is a significant health hazard^44^.

Some encountered representatives are used in human consumption. For example, mushrooms of the genera *Agaricus* are used directly for consumption, while *Penicillium* molds are used to ripen cheeses^46^, and *Saccharomyces* fungi are used in, biofuel, baking, and beer and wine industry^47^. *Beauveria bassiana* and several species of *Metarhizium* are entomopathogenic fungi that are used as biological insecticides^11,48^.

## 6. Bacterial taxa of interest

We encountered several bacteria that are potential agricultural pests as well as of medical importance to humans. For example, species of *Rickettsia* and *Wolbachia* are associated with a large variety of arthropod taxa, and are linked to both plant and human diseases^49,50^. We found several species of *Acinetobacter*, whose members are an important part of soil and water microbiomes^51^. Some species are human pathogens, and we encountered the potentially pathogenic *A. ursingii*^52^. Members of *Staphylococcus* are mostly harmless to humans, and occur in soil and on plant flowers^53^. Several species of *Xanthomonas* cause plant diseases^54,55^. *Salmonella* occurs in the digestive tract and skin of warm and coldblooded animals, and is medically important for humans^56^. *Salmonella* and *Clostridium*, also encountered in our samples, can be present in flies and beetles, especially when collected around livestock^57^. Some species of *Enterococcus* cause human infections^58^. While some members of *Haemophilus* are important human pathogens, many have a range of hosts, and many are commonly found in invertebrate intestines^56,59^. Other genera with plant pathogenic representatives include *Pseudomonas*, *Erwinia* that typically infect woody plants, and *Dickea* and *Pectobacterium* that are mostly pathogens of herbaceous plants^55^. *Pseudomonas pertucinogena* produces pertucin, a bacteriocin active against the whooping cough causing *Bordetella pertussis*^60^.

